# Divergent phenotypic plasticity of a convergent Mendelian trait in *Drosophila*

**DOI:** 10.1101/2021.11.10.468073

**Authors:** Pascaline Francelle, Jean R. David, Amir Yassin

## Abstract

In *Drosophila*, comparisons of the thermal plasticity of pigmentation across serially homologous abdominal segments have been conducted in two species, namely *Drosophila melanogaster* and *D. kikkawai*. Pigmentation variation has different genetic architecture in the two species, being oligogenic in the former and monogenic in the later. Here, we analyze the thermal plasticity of abdominal pigmentation in a third species, *D. erecta*, which is phylogenetically close to *D. melanogaster* but like *D. kikkawai* has a monogenic basis for pigmentation variation. However, the underlying locus differs between *D. erecta* and *D. kikkawai*, being the X-linked melanin-synthesis gene *tan* in the former and the autosomal transcription factor *pdm3* in the later. We found that in spite of a low overall plasticity in monogenic species compared to *D. melanogaster*, the two monogenic species showed divergent plasticity patterns in respect to the response to temperature and to the degree of dominance in heterozygotes. Those results provide new insights on the dependence of the degree of plasticity on the genetic architecture as well as on the extent of phenotypic convergence.

## Introduction

Phenotypic plasticity is the ability of a genotype to produce different phenotypes under different environmental conditions (Pigliucci 2001). Plasticity may have a major role in facilitating adaptation and promoting phenotypic evolution (West-Eberhard 2003). In spite of significant progress in unraveling the molecular mechanisms underpinning plasticity in several model organisms, patterns of plasticity evolution remain largely unclear (Sommer 2020). A particular aspect that has attracted lesser attention is the plasticity of the same trait on serially homologous anatomical structures, such as teeth, phalanges or body segments. Whereas *trans*- and *cis*-regulatory modification of genes of the same network could lead to phenotypic differences among structures, how the phenotype at each structure responds to environmental stimuli is still unclear. To the best of our knowledge, this question has mainly been addressed in a few traits, such as the wing eyespots in butterflies (Monteiro *et al.* 2015; Bhardwaj *et al.* 2020), and abdominal pigmentation in *Drosophila* (Gibert *et al.* 1998, 1999, 2000, 2008, 2016, 2017; Castro *et al.* 2018).

*Drosophila* pigmentation is a well-characterized genetic trait resulting from a conserved network of melanin synthesis genes, such as *yellow*, *tan* or *ebony*. Pigmentation differences between body segments, sexes, geographical populations and species are often due to *cis*- and *trans*-regulatory modifications of those melanin synthesis genes (Massey and Wittkopp 2016; Hughes *et al.* 2021). In *D. melanogaster*, cold developmental environments produce dark flies due to changes in the expression of *tan* (Gibert *et al.* 2016), *yellow* (Gibert *et al.* 2017) and *bric-à-brac* (Castro *et al.* 2018). Thermal plasticity, however, differs in this species between body segments and the sexes (Gibert *et al.* 2000). In females, the phenotypic plasticity between cold and warm environments increases towards most posterior abdominal segments, with the pigmentation of the seventh abdominal segment (P7) being the most plastic. In males, on the other hand, this pattern is restricted to the first three abdominal segments (P2, P3 and P4), whereas the last two segments (P5 and P6) are completely dark with almost no plastic response.

The pattern of pigmentation plasticity across abdominal segments was, however, investigated in a few other *Drosophila* species, most notably in *D. kikkawai* (Gibert *et al.* 1999). This species belongs the *montium* species group and shared a last common ancestor with *D. melanogaster* more than 20 million years ago (Conner *et al.* 2021). Like several members of the *montium* group, *D. kikkawai* is characterized by the presence of a female-limited color dimorphism (FLCD) that is caused by a single autosomal locus (Freire-Maia 1949; Ohnishi and Watanabe 1985). In four species including *D. kikkawai*, the causative locus of this Mendelian FLCD mapped to the transcription factor *pdm3* (Yassin *et al.* 2016b). The overall level of abdominal pigmentation plasticity between *D. kikkawai* and *D. melanogaster* differs, with the latter being more plastic. It is, however, unknown whether the difference between the two species is due to their long divergence, leading to accumulative changes in the melanin synthesis genes, or to the evolution of the genetic architecture of pigmentation variation being monogenic in *D. kikkawai* and oligogenic in *D. melanogaster*.

To test whether divergence in pigmentation plasticity across serially homologous abdominal segments between *D. melanogaster* and *D. kikkawai* was due to phylogenetic divergence or changes in the number of underlying genes, we analyzed here this trait in a third species, *D. erecta*. *D. erecta* is a member of the *melanogaster* species subgroup, and consequently it is closely-related to *D. melanogaster*. However, unlike most species of the *melanogaster* subgroup, *D. erecta* is unique in having evolved a Mendelian FLCD reminiscent to that of the *montium* species. Remarkably, the FLCD locus in *D. erecta* is X-linked (Payant 1988) and it maps to a short regulatory sequence of the melanin-synthesis gene *tan* (Yassin *et al.* 2016a). We found that in spite of the phenotypic convergence of FLCD in *D. kikkawai* and *D. erecta*, the two species showed distinct thermal plasticity responses in relation to the sexes and the body segments. *D. erecta* thermal responses were also distinct from those of *D. melanogaster*, strongly supporting an important effect of the genetic architecture of a trait on the evolution of its phenotypic plasticity.

## Materials and Methods

We analyzed the same *D. erecta* parental strains used by Yassin *et al.* (2016a), *i.e.* C3 and NN, which are homozygous for the light and dark FLCD allele, respectively. The C3 strain was established from a Gabonese population collected from Gabon in 2006. The NN strain was obtained by introgressing the dark allele from the same population in the C3 strain for 12 backcrossed generations before establishing a homozygous dark line. This line differs from the parental C3 light parental line in only 1 Mb introgressed region centered on the *cis*-regulatory element of the *tan* region on chromosome X.

For each parental homozygous strain, groups of 10 males and 10 females were placed in separate vials containing standard rearing medium. To obtain heterozygous flies, reciprocal crosses involving 10 males and 10 females each sex from a different parental line were also placed in separate vials. The vials containing the offspring of all crosses were placed in separate incubators at six growth temperatures: 14, 16, 21, 25, 28 and 30°C. were were and their heterozygous F1 offspring resulting from reciprocal crosses. Pigmentation was analysed on 10 flies from each sex from each parental strain and the two F1 strains. Pigmentation was scored on successive tergites (P2 to P7 in females and P2 to P6 in males) according to the surface of black area at the posterior part of each tergite by establishing 11 phenotypic classes from 0 (no black pigment) up to 10 (tergite completely black) as in David *et al.* (1990). Pigmentation scores on successive abdominal segments in different growth temperature were obtained from the literature for *D. kikkawai* (Gibert *et al.* 1999) and for a strain of *D. melanogaster* from Bordeaux, France (Gibert *et al.* 2000). All statistical analyses were conducted using R (R Core Team 2016), with 3d plots obtained using the Plotly package in R (Sievert 2020).

## Results

Patterns of pigmentation plasticity in both females and males in respect to abdominal segment and developmental temperature are summarized in Figures 1 and 2, respectively. For all species, a similar pattern of darker flies under colder conditions can be observed. However, the relative significance of each of the independent factors greatly differs between the three species (Table 1). In *D. erecta*, body segmentation accounts for ~44% of the variance and it is the only factor with a variance explanation >10%. In *D. melanogaster*, body segmentation is also the most explanatory factor (~37%), but several other factors have effects of >10%, *i.e.* temperature, sex and sex x segmentation. In *D. kikkawai*, sex is the most important factor (~29% of the variance) followed by segmentation, genotype and their interaction.

**Figure 1.**
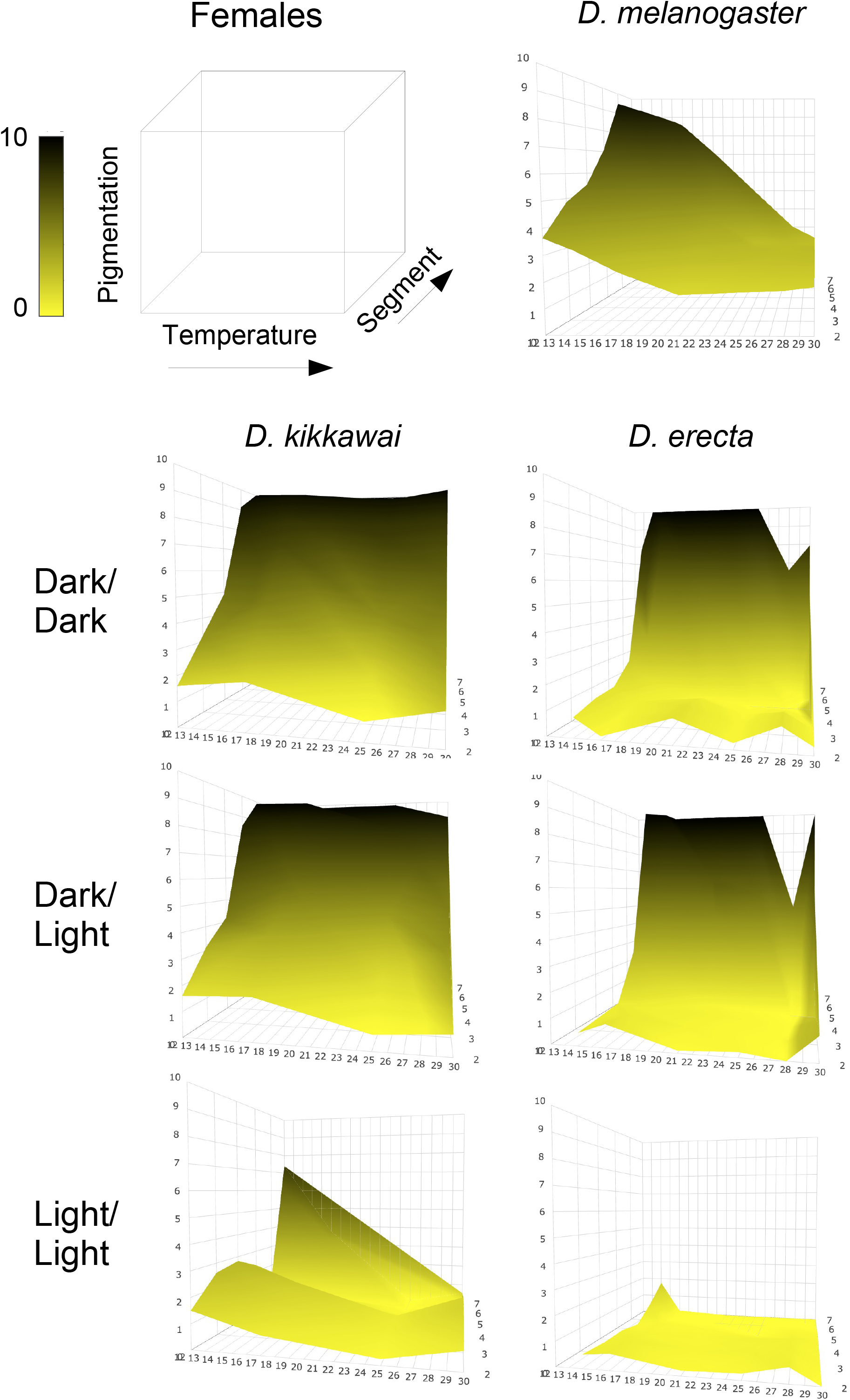
3D-mesh plot of thermal phenotypic plasticity of pigmentation on successive abdominal segments in females of three *Drosophila* species.

**Figure 2.**
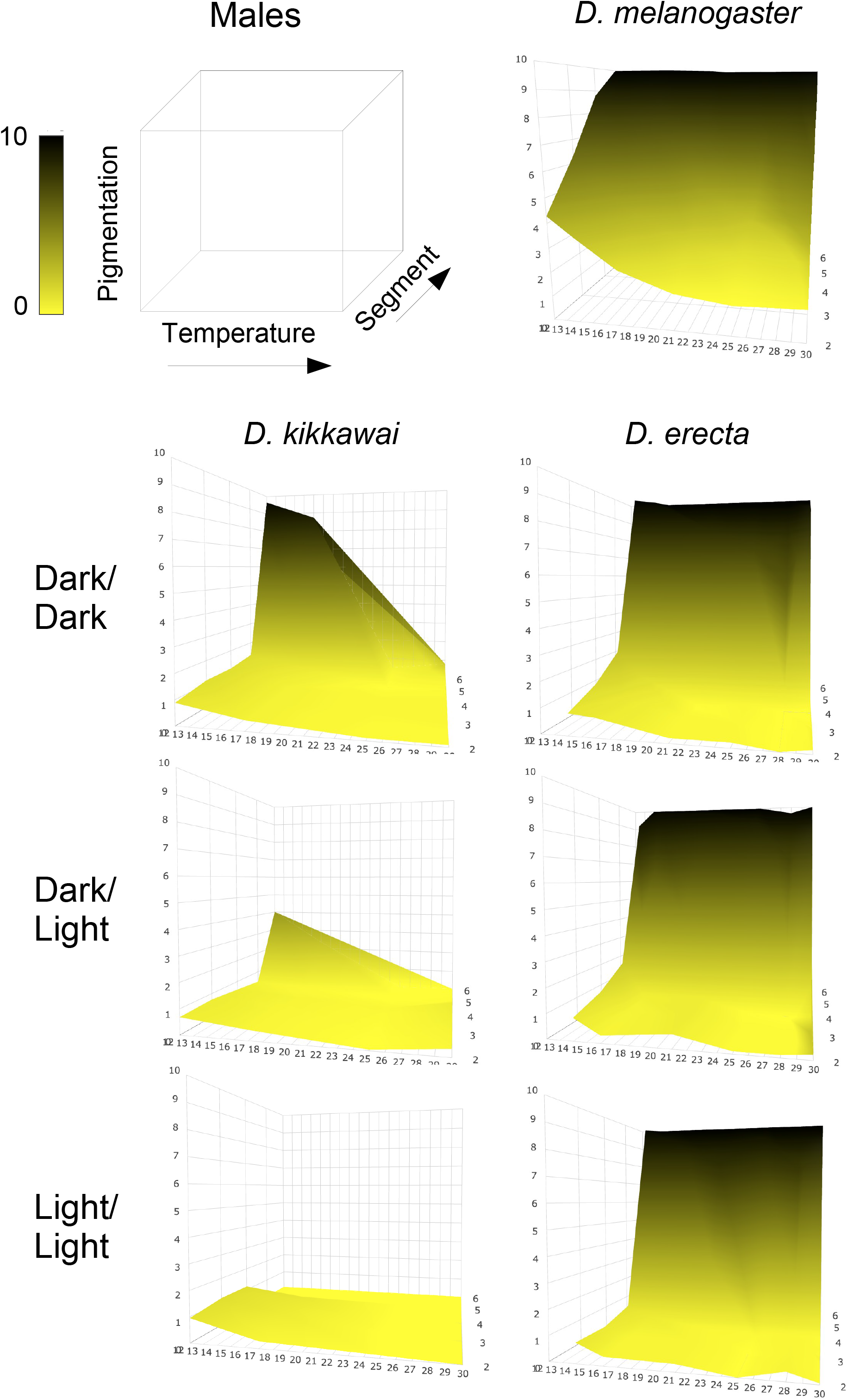
3D-mesh plot of thermal phenotypic plasticity of pigmentation on successive abdominal segments in males of three *Drosophila* species.

**Table 1.** Proportions of variance (%) explained by each sex, genotype, abdominal segment and their interactions as deduced from multi-factorial Analysis of Varience (ANOVA) in *D. melanogaster*, *D. erecta* and *D. kikkawai*. ANOVA significance levels are given as: * <0.05, ** <0.01 and *** <0.001. Note that no genotype was analysed for *D. melanogaster* since this species is oligogenic.

| Factor | *D. melanogaster* | *D. erecta* | *D. kikkawai* |
| --- | --- | --- | --- |
| Sex | 10.77*** | 5.38*** | 28.69*** |
| Genotype |  | 3.44*** | 12.27*** |
| Segment | 37.28*** | 43.97*** | 16.76*** |
| Temperature | 19.69*** | 2.73*** | 2.63*** |
| Sex x Genotype |  | 2.47*** | 6.63*** |
| Sex x Segment | 13.22*** | 0.26 | 8.22*** |
| Genotype x Segment |  | 3.45*** | 10.33*** |
| Sex x Temperature | 1.77*** | 0.09 | 0.32 |
| Genotype x Temperature |  | 0.33 | 0.47 |
| Segment x Temperature | 2.03*** | 0.95** | 1.01** |
| Sex x Genotype x Segment |  | 3.05*** | 4.72*** |
| Sex x Genotype x Temperature |  | 0.15 | 0.82** |
| Sex x Segment x Temperature | 5.32*** | 0.11 | 1.06* |
| Genotype x Segment x Temperature |  | 0.13 | 1.07 |
| Sex x Genotype x Segment x Temperature |  | 0.13 | 1.40** |
| Residuals | 9.93 | 33.35 | 3.60 |

The degree of plasticity, as measured by the average phenotypic variance across environments, differed between the three species (Table 2). *D. melanogaster* females and males were the most plastic (*σ*^*2*^= 3.83 averaged over the two sexes, Figures 1 and 2), followed by *D. erecta* (*σ*^*2*^= 1.46) then *D. kikkawai* (*σ*^*2*^= 1.22). For *D. erecta*, homozygous dark females and males were the most plastic, whereas in *D. kikkawai*, opposite genotypes between the sexes were the most plastic, the homozygous light in females and the homozygous dark in males. The abdominal segment with the highest phenotypic variance across environments also differed between species and sexes. In *D. melanogaster* it was P7 (σ^2^ = 18.18) and P4 (σ^2^ = 5.81) for females and males, respectively. In *D. erecta*, those were P6 (σ^2^ = 16.79) and P5 (σ^2^ = 2.98) in homozygous dark females and males, respectively. In *D. kikkawai*, the most plastic segments were P7 in homozygous light females (σ^2^ = 9.63) and P6 in homozygous dark males (σ^2^ = 16.72).

**Table 2.** Degree of plasticity in terms of phenotypic variance (*σ*^*2*^) of pigmentation scores of successive abdominal segments across different growth temperatures.

| Species | Sex | Strain | P2 | P3 | P4 | P5 | P6 | P7 |
| --- | --- | --- | --- | --- | --- | --- | --- | --- |
| *melanogaster* | Female | Bordeaux | 0.69 | 1.03 | 1.52 | 3.70 | 8.43 | 18.18 |
| *erecta* | Female | C3 | 0.07 | 0.02 | 0.00 | 0.02 | 0.13 | 0.50 |
| *erecta* | Female | F1 | 0.14 | 0.16 | 0.20 | 0.64 | 14.41 | 1.78 |
| *erecta* | Female | NN | 0.17 | 0.11 | 0.05 | 1.40 | 16.79 | 4.17 |
| *kikkawai* | Female | LL | 0.16 | 0.38 | 0.38 | 0.44 | 0.29 | 9.63 |
| *kikkawai* | Female | F1 | 0.12 | 0.37 | 0.86 | 2.16 | 1.12 | 0.05 |
| *kikkawai* | Female | DD | 0.15 | 0.57 | 0.92 | 0.83 | 0.03 | 0.00 |
| *melanogaster* | Male | Bordeaux | 1.48 | 2.88 | 5.89 | 0.04 | 0.00 |  |
| *erecta* | Male | C3 | 0.08 | 0.07 | 0.15 | 0.78 | 0.00 |  |
| *erecta* | Male | F1 | 0.18 | 0.31 | 0.76 | 0.99 | 0.01 |  |
| *erecta* | Male | NN | 0.12 | 0.31 | 0.98 | 2.98 | 0.00 |  |
| *kikkawai* | Male | LL | 0.10 | 0.17 | 0.14 | 0.00 | 0.00 |  |
| *kikkawai* | Male | F1 | 0.05 | 0.08 | 0.07 | 0.01 | 3.45 |  |
| *kikkawai* | Male | DD | 0.07 | 0.12 | 0.11 | 0.24 | 16.72 |  |

The monogenic nature of FLCD variation in *D. erecta* and *D. kikkawai* allowed to compare plasticity in genetic dominance in heterozygotes issued from reciprocal crosses in each species (Figure 3). Whereas dominance reversal in the last abdominal segments between sexes has previously been found in *D. kikkawai*, no such dominance reversal between the sexes was found in *D. erecta*, where males are hemizygous for the X-linked FLCD locus.

**Figure 3.**
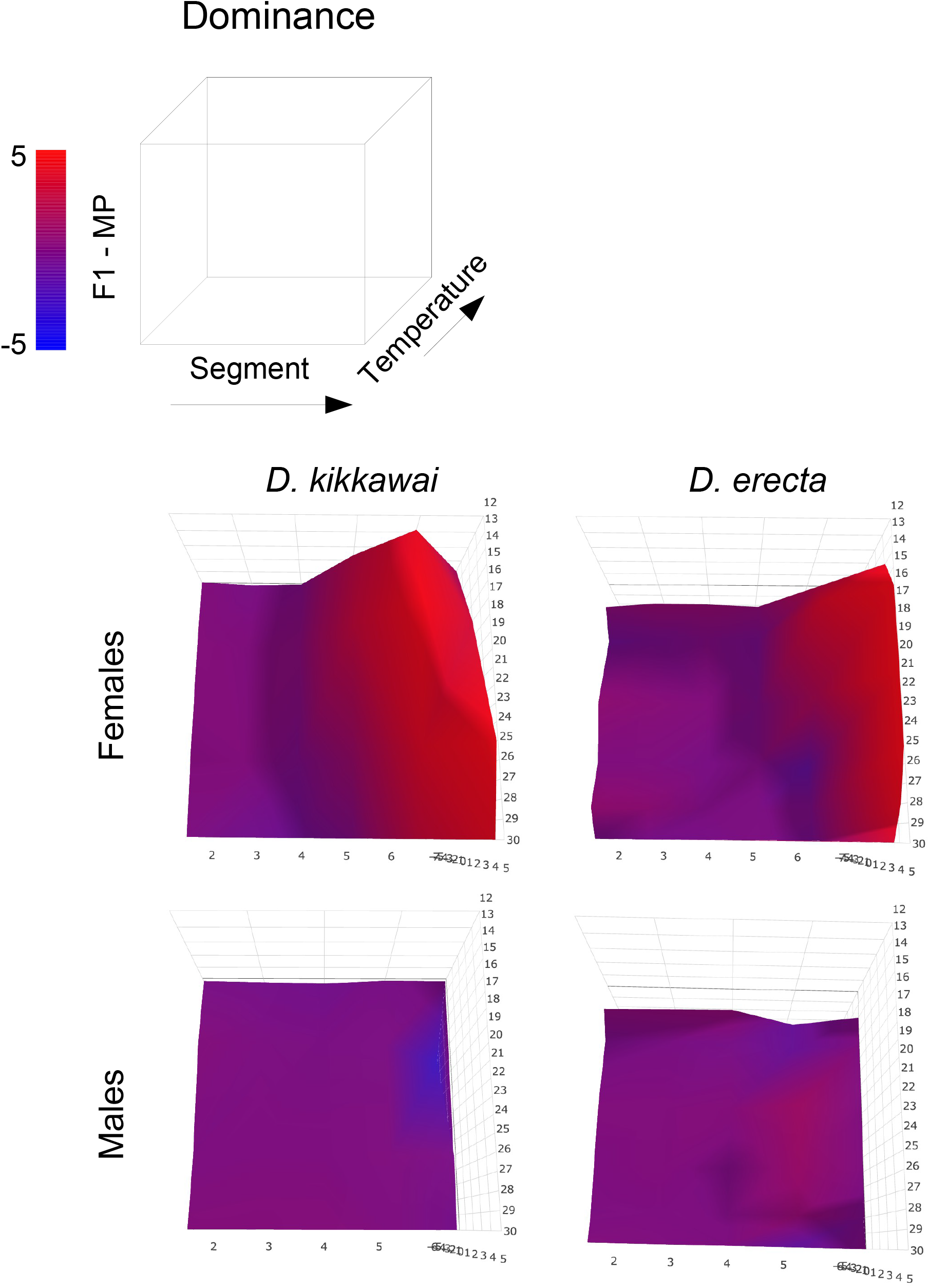
3D-mesh plot of thermal phenotypic plasticity of pigmentation dominance on successive abdominal segments in heterozygous females and males of *Drosophila kikkawai* and *D. erecta*.

## Discussion

Taken together, our results provide insights on the genetic and developmental basis of pigmentation plasticity evolution in *Drosophila*. Pigmentation plasticity may depend on the number of underlying loci. This is most obvious in comparison between *D. melanogaster* and *D. erecta*, both species belonging to the *melanogaster* subgroup. In *D. melanogaster* females, wherein pigmentation variation has an oligogenic architecture (Bastide *et al.* 2013, 2016; Dembeck *et al.* 2015), all segments are plastic with plasticity tending to increase with decreasing temperature and toward posterior segments. In the monogenic *D. erecta* (Yassin *et al.* 2016a), females show no plastic changes in response to temperature regardless to the underlying genotype. Indeed, females lack of response in this species resembles that of conspecific males as well as males of *D. melanogaster*. For the monogenic *D. kikkawai* (Yassin *et al.* 2016b), which is phylogenetically distant from both species, plasticity in a way similar to that of *D. melanogaster* is found for homozygous light females and homozygous dark males. However, for homozygous dark females, homozygous light males and heterozygous males and females, pigmentation shows no plastic response to growth temperature.

In spite of the phenotypic convergence between *D. erecta* and *D. kikkawai*, FLCD in both species shows contrasting thermal plasticity. This observation significantly bears on current debates about the limitedness of morphological evolution. Indeed, several authors have suggested that morphological convergence may be so ubiquitous in evolution that novelties are rare (McGhee 2011; Blount *et al.* 2018; Pigot *et al.* 2020). A phylogenetic analysis of 490 morphological characters in *Drosophila* suggested that for every morphological change there are twice the chances to derive a homoplasic character state than to derive a novelty (Al Sayad and Yassin 2019). However and for multiple practical reasons, evolutionary biologists rarely investigate phenotypic plasticity in defining their traits in phylogenetic analyses. The study presented here is a reminder that even in the most obvious cases of phenotypic similarities between distant species, phenotypic plasticity may show innovative patterns that would be overlooked when traits are compared under single environments. Future genetic work should help elucidating how *tan* and *pdm3* expression differs between thermal regimes in *D. erecta* and *D. kikkawai*, and how much their difference in expression alters other components of the melanin-synthesis pathway, leading to the divergent plasticity we documented here.

## Conflicts of interest

The authors declare no conflict of interest.

## Notes

### Competing Interest Statement

The authors have declared no competing interest.

